# *Lacticaseibacillus rhamnosus* GR-1 attenuates *Escherichia coli*-induced endometritis in mice, accompanied by modulation of uterine microbiota and metabolite profiles

**DOI:** 10.64898/2026.02.09.704851

**Authors:** Xiaohan Li, Li Jia, Mengsen Wang, Xiangfu Wen, Deyuan Song, Guowei Liang, Dengke Liu, Mingchao Liu

## Abstract

**Objective:** Endometritis in dairy cows is a highly prevalent infectious disease in cattle farming, which can cause persistent inflammatory responses in the uterus. This study aimed to evaluate the therapeutic efficacy of *Lacticaseibacillus rhamnosus* (*L. rhamnosus*) GR-1 against *Escherichia coli* (*E. coli*)-induced endometritis in mice and investigate its regulatory effects on uterine microbiota and metabolites.

**Methods:** A total of 36 female BALB/c mice aged 7-8 weeks were randomly divided into 4 groups: CONT group, ECOL group (1.0×10^8^ CFU/mL *E. coli*), AMP group (0.1 mg/kg ampicillin), and GR-1 group (1.0×10^9^ CFU/mL *L. rhamnosus* GR-1). Uterine morphological characteristics were observed, and the expression of uterine inflammatory cytokines, oxidative apoptosis-related genes and proteins, as well as related reproductive indices were detected. 16S rRNA sequencing and metabolomic sequencing were performed to evaluate the profiles of uterine microbiota and metabolites.

**Results:** *L. rhamnosus* GR-1 effectively alleviated *E. coli*-induced uterine edema and inflammatory infiltration, and modulated the levels of inflammatory factors (MPO, IL-6, IL-1β, TNF-α, IL-10 and MDA) in the uterus. Meanwhile, this strain elevated the levels of antioxidant factors (GSH-Px and SOD), inhibited the expression of the oxidative apoptosis-related factors Keap1, Bax and caspase-3, and upregulated that of Nrf2, HO-1, NQO1 and Bcl-2. Furthermore, treatment with *L. rhamnosus* GR-1 increased the abundance of beneficial uterine microbiota, promoted the accumulation of L-isoleucyl-L-proline and N-acetylcysteine, and ameliorated the pregnancy status of mice.

**Conclusion:** *L. rhamnosus* GR-1 enhanced uterine anti-inflammatory and antioxidant stress capacity to alleviate *E. coli*-induced endometritis in mice. It modulated uterine microbiota and metabolites, and also improved the reproductive capacity of mice. Thus it may provide a potential antibiotic alternative for the treatment of dairy cow endometritis.

## INTRODUCTION

As one of the most common diseases in the dairy cattle industry, endometritis leads to decreased milk production and excessive uterine inflammatory responses, which severely threatens the development of the dairy cattle industry and causes substantial economic losses to the breeding sector [1]. Clinically, postpartum endometritis leads to a 10-20% decline in milk production per affected cow and a significant increase in culling rates due to reproductive failure [2]. The pathogenesis of dairy cow endometritis is primarily driven by dysregulated interactions among pathogenic infection, uterine microbiota imbalance, and impaired mucosal immunity, with *Escherichia coli* (*E. coli*) identified as the principal etiological agent [3]. During the periparturient period of dairy cows, *E. coli* tends to proliferate rapidly in the uterine cavity. This excessive proliferation triggers the release of virulence factors such as lipopolysaccharide (LPS) and hemolysin, disrupting the integrity of the uterine mucosal barrier, inducing excessive inflammatory responses, and causing uterine tissue damage [4,5].

Current clinical management of *E. coli*-induced endometritis in dairy cows primarily relies on broad-spectrum antibiotics such as ampicillin [6]. But the overuse of antibiotics exacerbates antimicrobial resistance (AMR) [7]. Some studies have shown that the *E. coli* detected in endometritis-affected dairy cows are mainly multidrug-resistant (MDR) strains [8]. This not only increases the risk of recurrent infections and higher culling rates of dairy cows but also facilitates the transmission of resistance genes, posing a potential threat to public health. Additionally, while antibiotics kill pathogenic bacteria, they also eliminate certain *Lactobacillus* strains that produce lactic acid and bacteriocins. The loss of this microbial community further increases the risk of secondary infections and chronic inflammation in dairy cows [9].

Probiotics are a class of active microbes that can establish colonization on the mucosal tissues of intestinal and reproductive systems in hosts [10]. *L. rhamnosus* GR-1 is a strain with well-documented probiotic potential, which has been proven to regulate mucosal immunity and restore microbiota balance in the animal reproductive tract [11,12]. Previous studies confirm that *L. rhamnosus* GR-1 promotes antioxidant factor secretion and coordinates apoptotic factor release to maintain mitochondrial function, thereby exerting a protective effect on bovine endometrial epithelial cells [13]. However, the efficacy of *L. rhamnosus* GR-1 against *E. coli*-induced endometritis remains unclear. However, the effect of *L. rhamnosus* GR-1 on *E. coli*-induced endometritis remains unclear. Therefore, this study investigated the therapeutic effects of *L. rhamnosus* GR-1 on *E. coli*-induced endometritis.

## MATERIALS AND METHODS

### Bacteria strains and reagents

*Lacticaseibacillus rhamnosus* GR-1 (ATCC 55826) was supplied by the Laboratory of Clinical Nutrition and Immunology, College of Veterinary Medicine, China Agricultural University. The isolate was grown in MRS medium (AOBOX, Beijing, China) at 37°C, and the culture was subsequently resuspended in PBS to obtain a final concentration of 1.0×10⁸ CFU/mL for later use. *Escherichia coli* O111:K58 (CVCC1450) was obtained from the same laboratory. This *E. coli* strain was incubated in Luria–Bertani (LB) broth (AOBOX) at 37°C with shaking at 180 rpm, after which it was diluted in PBS to 1.0×10⁸ CFU/mL to establish the mouse uterine inflammation model. Ampicillin (Cat. No. A6920; purity >96%) (Solarbio, Beijing, China) was used as the positive control drug.

### Animals and experiment design

All procedures involving mice were approved by the Animal Care and Use Committee of Hebei Agricultural University (Approval Number: 2021041) and were conducted in accordance with the approved guidelines. Thirty-six female BALB/c mice (Spiceff, Beijing, China) aged 7-8 weeks with a body weight of 17±2 g were maintained under controlled temperature and humidity conditions (24 ± 1°C; 40%–80% relative humidity) with ad libitum access to chow and water. Before the formal trial, the working dose of *Lactobacillus rhamnosus* GR-1 was optimized; 1.0 × 10⁹ CFU/mL was selected for subsequent experiments (Supplement 4). After 1 week of acclimatization, the 36 mice were randomly and equally allocated to four groups (n = 9 per group): Control (CONT), infected control (ECOL), ampicillin control (AMP), and *L. rhamnosus* GR-1 treatment (GR-1). Days 0–2: the CONT group received uterine horn perfusion with 100 μL PBS, whereas mice in the ECOL, GR-1, and AMP groups were administered 100 μL of *Escherichia coli* suspension (1.0 × 10⁸ CFU/mL). Days 3–6: uterine horn perfusion with 100 μL PBS was continued for the CONT and ECOL groups; the GR-1 group was perfused with 100 μL *L. rhamnosus* GR-1 (1.0 × 10⁹ CFU/mL); and the AMP group received 100 μL containing 0.1 mg ampicillin. On Day 7, six mice from each group were randomly chosen, euthanized, and uterine tissues were harvested. From Days 7–12, the remaining animals in each group (n = 3) were kept with free access to food and water. On Day 12, one male mouse was randomly introduced into each group for mating, and males were removed on Day 19. Reproductive outcomes were monitored until parturition occurred in females (Figure 1A). In addition, after body weight was recorded, an experimental animal thermometer (ZK-FT3400; Zhike Hongrun Environmental Protection, Henan, China) was used to determine body temperature changes in each group, Determination of MPO (Solarbio) content in mouse uterine tissues.

**Figure 1.**
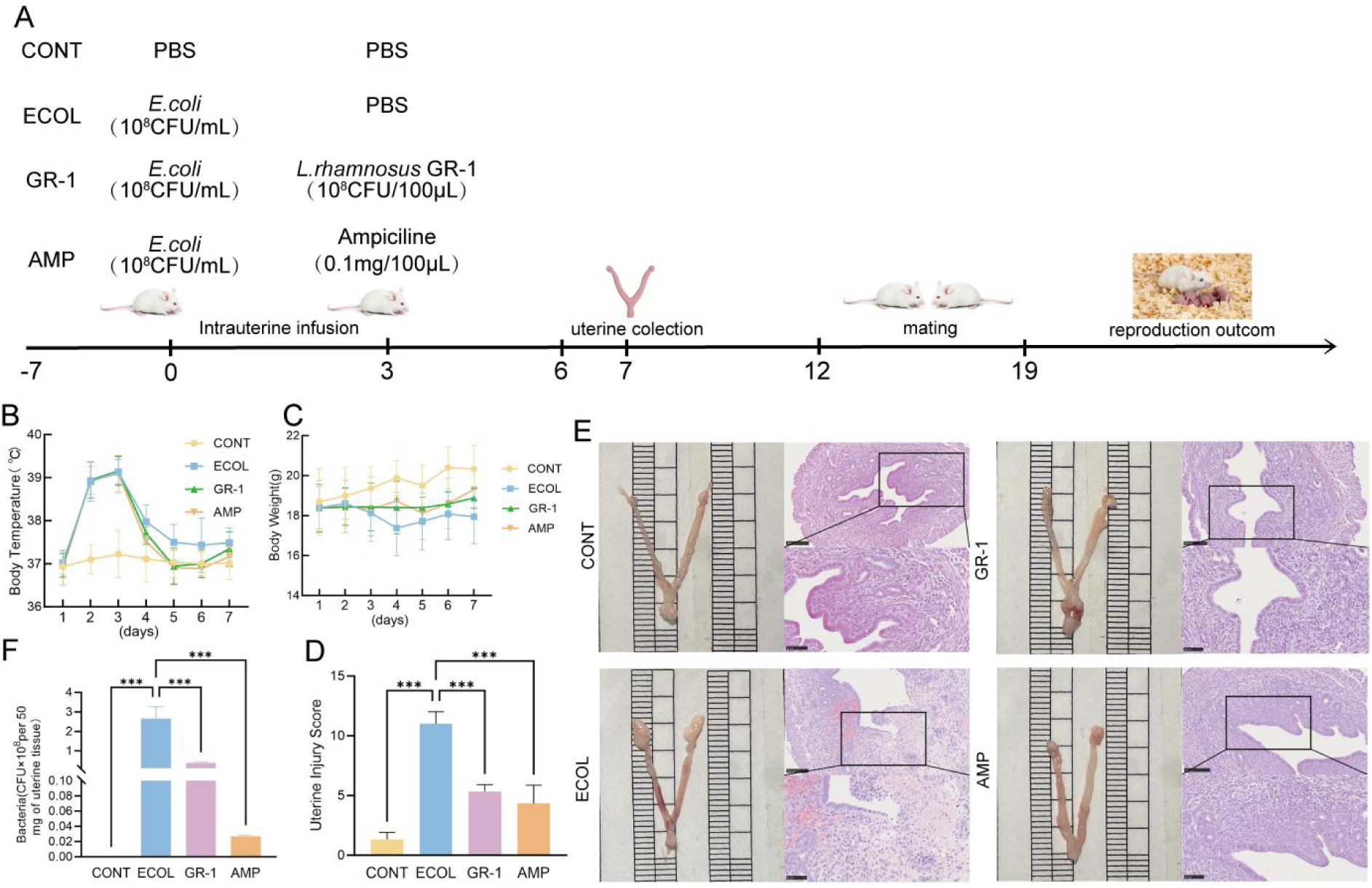
Effects of *L. rhamnosus* GR-1 on *E.coli*-induced endometritis. (A) Mouse experimental design. Four groups were set up: From day 0 to day 2, except for the CONT group, the uteri of mice in the other groups were infused with *E.coli* at a concentration of 1.0×10⁸ CFU/mL. From day 3 to day 6, the CONT group and ECOL group were given PBS; the GR-1 group was given *L. rhamnosus* GR-1 (1.0×10⁹ CFU/mL, 100 μL); the AMP group was given ampicillin (0.1 mg/100 μL). On day 7, some mice were sacrificed. From day 7 to day 12, the “mating and breeding period” was conducted. Group abbreviations: CONT (control group), ECOL (*E.coli* group), GR-1 (*L. rhamnosus* group), AMP (ampicillin group),. (B) Changes in body temperature, n=6. (C) Changes in mouse body weight, n=6. (D) Analysis of histological scores. (E) Histomorphology of the uterus, n=3. (F) Uterine tissue bacterial, n=6. Data represented are mean ± SD, *p<0.05; **p<0.01; ***p<0.001.

### Uterine bacterial load

Fifty milligrams of uterine tissue were harvested, mixed with grinding beads, homogenized, and then serially diluted. Next, 100 μL of the resulting bacterial suspension was plated on eosin methylene blue (EMB) agar (AOBOX), while another 100 μL was inoculated into LB broth for culture. After incubation, the *E. coli* burden in each group was determined by colony enumeration using standard plate-count methods.

### Histopathological evaluation

Uterine tissues from mice were collected and preserved in 4% paraformaldehyde for 48 h, and then processed for paraffin embedding. Sections were cut from the paraffin blocks at 5 μm thickness and stained with hematoxylin and eosin (H&E). The prepared slides were subsequently examined and assessed using a fluorescence microscope (IX51; Olympus, Tokyo, Japan). The specific scoring standards are described in Supplement 1.

### Uterine oxygen-related factor levels

According to the manufacturer’s (Solarbio) instructions, the levels of glutathione peroxidase (GSH-Px), total superoxide dismutase (SOD), and malondialdehyde (MDA) in uterine tissues of mice in each experimental group were detected. Optical density was measured using a microplate spectrophotometer (Synergy LX, BioTek).

### Enzyme-Linked Immunosorbent Assay for uterine inflammatory mediators

The inflammatory-related factors in mouse uterine tissues, including tumor necrosis factor-α (TNF-α), interleukin-1β (IL-1β), interleukin-10 (IL-10), and interleukin-6 (IL-6), were detected using a kit (Jingmei, Jiangsu, China) and measured by a microplate spectrophotometer (Synergy LX) for optical density.

### Quantitative real-time PCR (RT-qPCR)

To quantify the mRNA expression of heme oxygenase-1 (HO-1), nuclear factor erythroid 2–related factor 2 (Nrf2), Kelch-like ECH-associated protein 1 (Keap1), NAD(P)H quinone dehydrogenase 1 (NQO1), cysteinyl aspartate–specific proteinase 3 (Caspase3), BCL2-associated X protein (Bax), and B-cell lymphoma 2 (Bcl-2). Total RNA was isolated from mouse uterine tissues using TransZol Up Plus RNA Kit (TransGen, Beijing, China). Primer information is listed in Supplement 2. Amplification was performed using ChamQ Universal SYBR qPCR Master Mix (Vazyme, Nanjing, China) under the following conditions: an initial denaturation at 95°C for 30 s, followed by 40 cycles of denaturation at 95°C for 10 s and annealing/extension at 60°C for 30 s. Glyceraldehyde-3-phosphate dehydrogenase (GAPDH) served as the internal control. Relative gene expression was determined using the 2^−ΔΔCt approach.

### Transcriptional expression of genes related to oxidative stress

Uterine tissues were lysed in RIPA buffer (Solarbio) to extract total protein. Protein concentrations were determined using a BCA Protein Assay Kit (Solarbio). For SDS–PAGE, equal amounts of protein (10 μg per sample) were loaded and electrophoretically separated, and the resolved proteins were then transferred to PVDF membranes for 2 h. Membranes were blocked with a rapid blocking solution (Yamei, Beijing, China) for 20 min at room temperature, followed by incubation with the corresponding primary antibodies at 4°C overnight (antibodies are listed in Table 1). After washing, membranes were incubated with HRP-conjugated goat anti-rabbit IgG (H&L) for 1 h at room temperature. The blots were washed with TBST containing 0.1% Tween 20 (Solarbio), developed using an enhanced chemiluminescence (ECL) substrate (Xinsaimai, Jiangsu, China), and imaged with a chemiluminescence detection system (Odyssey, USA). Band intensities were quantified with ImageJ software.

**Table 1.**
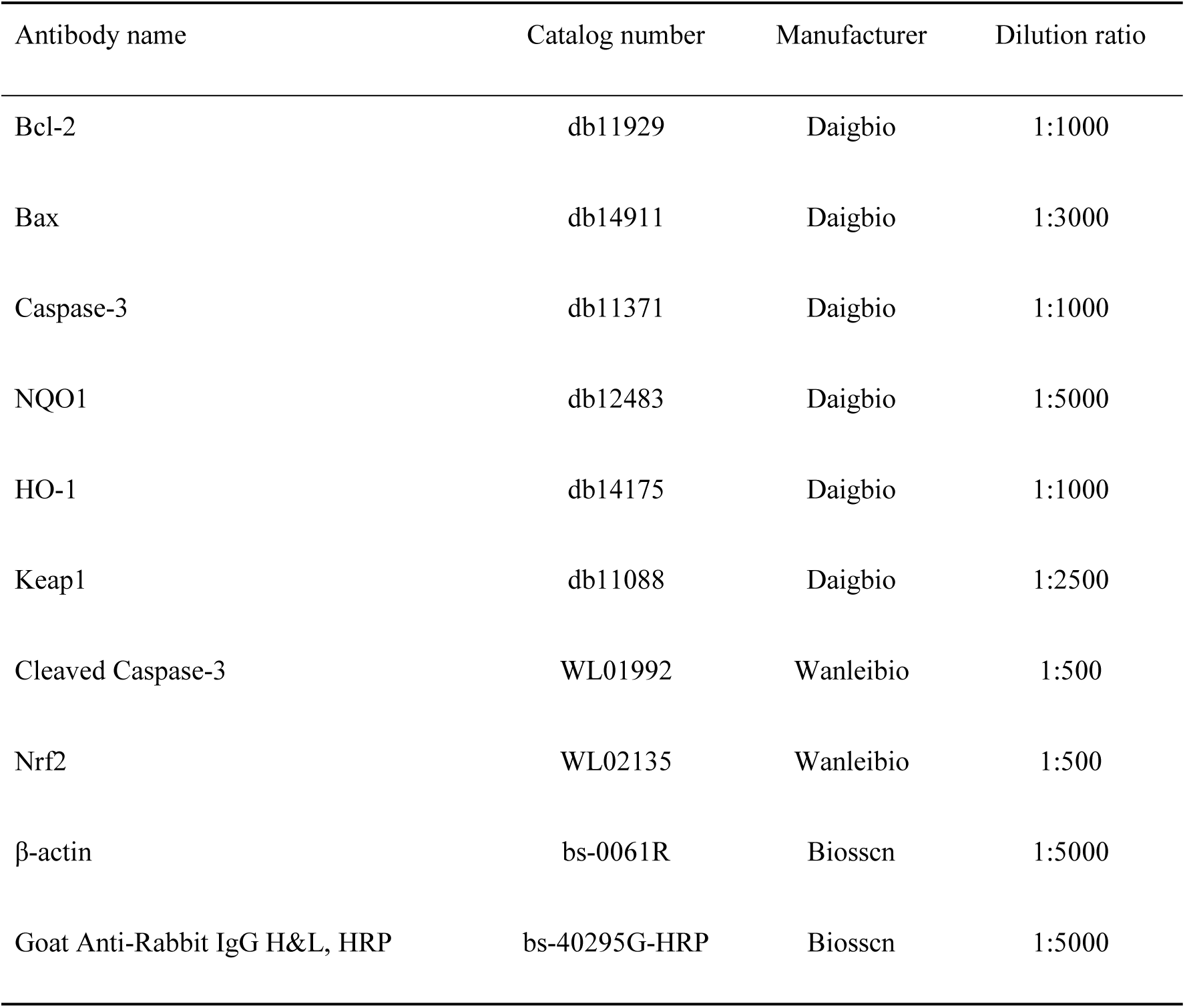
Antibodies used in western blot.

### Uterine 16S rRNA sequencing

Following pretreatment of the uterine samples, total DNA was isolated using the cetyltrimethylammonium bromide (CTAB) extraction procedure. Genomic DNA concentration was measured with a NanoDrop 2000 spectrophotometer (Thermo, Waltham, MA, USA), and DNA integrity was assessed by electrophoresis on a 1% agarose gel. The V3–V4 hypervariable region of the bacterial 16S rRNA gene was amplified by PCR with primers 515F and 806R, using a Bio-Rad T100 Gradient PCR Thermal Cycler (Model T100; Bio-Rad, Hercules, CA, USA). Amplicons were excised from 2% agarose gels and purified with a Universal DNA Gel Extraction Kit (Tiangen, Beijing, China). The purified products were then quantified on a Qubit® 2.0 Fluorometer (Thermo). Sequencing libraries were prepared according to the standard workflow with the NEBNext Ultra II DNA Library Prep Kit (New England Biolabs, Beijing, China). Finally, paired-end sequencing (PE250) was carried out on an Illumina NovaSeq 6000 platform following the manufacturer’s guidelines.

### Microbial metabolome sequencing and analysis in uterus

The uterine metabolomic profile was acquired on a Q Exactive™ HF/Q Exactive™ HF-X mass spectrometer interfaced with a Vanquish UHPLC system (Thermo). After data conversion and preprocessing, Partial Least Squares–Discriminant Analysis (PLS-DA) was applied to explore metabolic separation and identify intergroup differences. For pairwise comparisons, metabolite significance was evaluated using a t-test, and the fold change (FC) between groups was computed to reflect the magnitude of variation. Volcano plots were created in R with the ggplot2 package by jointly considering the Variable Importance in Projection (VIP) score, log₂(FC), and −log₁₀(p-value) to screen key metabolites; bubble plots were also generated using ggplot2. Metabolites were annotated against the KEGG database, and pathways with p<0.05 were defined as significantly enriched.

### Statistical analysis

Statistical analyses were conducted with GraphPad Prism 10 (San Diego, CA, USA), and data are expressed as mean ± standard deviation (SD). Normality was assessed before inferential testing. When datasets met normality assumptions, comparisons were performed using one-way ANOVA followed by Tukey’s multiple-comparisons test. If the data were not normally distributed, Dunnett’s T3 procedure was used. Statistical significance was defined as p<0.05. Analyses of uterine microbiota datasets were carried out using the Novogene Cloud Platform (https://cloud.majorbio.com). Associations between the microbiome and metabolome were evaluated with Spearman’s rank correlation; correlations were classified as strong when |r| ≥ 0.7, moderate for 0.5 ≤ |r| < 0.7, and weak when |r| < 0.5. Additional analytical details are provided in the figure legends for each panel.

## RESULTS

### Effects of *L. rhamnosus* GR-1 on the uterine morphology of mice infected with *E. coli*

After intrauterine exposure to *E. coli* (days 0–2), mice showed a decrease in body weight and an increase in body temperature. Following treatment with *L. rhamnosus* GR-1 or ampicillin during days 3–6, body weight rose and body temperature returned (Figure 1B–1C). As presented in Figure 1D–1E, uterine histoarchitecture in the CONT group remained intact and normal. The ECOL group exhibited obvious uterine edema, prominent inflammatory-cell infiltration, and a markedly higher histopathological abnormality score than the CONT group (p<0.001), averaging about 11 points. Administration of GR-1 orAMP reduced the score to approximately 5–6 points (p<0.001). In addition, Figure 1F shows that the uterine *E. coli* colony-forming units (CFU) were substantially increased in the ECOL group (p<0.001). The uterine CFUs was significantly reduced in mice treated with *L. rhamnosus* or ampicillin (p<0.001).

### Effects of *L. rhamnosus* GR-1 on uterine inflammation and oxidative stress induced by *E. coli* in mice

ELISA measurements of inflammatory mediators indicated that *E. coli* challenge markedly elevated TNF-α, IL-1β, and IL-6 levels in mouse uterine tissues (p<0.001), while IL-10 was significantly decreased (p<0.01). Notably, administration of *L. rhamnosus* GR-1 suppressed the *E. coli*–induced increases in TNF-α, IL-1β, and IL-6, and concurrently enhanced IL-10 production (p<0.01) (Figure 2A). *E. coli* exposure significantly increased uterine MPO and MDA activities (p<0.001), while causing a significant decrease in SOD and GSH-Px activities (p<0.001).The *L. rhamnosu*s treatment reduced uterine MPO and MDA activities (p<0.001) and significantly elevated SOD and GSH-Px activities (p<0.001) (Figure 2B). After *E. coli* stimulation, Keap1 mRNA expression was upregulated (p<0.001), whereas HO-1, NQO1, and Nrf2 transcripts were downregulated. The *L. rhamnosu*s significantly decreased Keap1 expression (p<0.001) and markedly increased HO-1, Nrf2, and NQO1 expression (Figure 3E). Regarding apoptosis, *E. coli* exposure significantly increased mRNA levels of the Bax and Caspase-3 (p<0.001), while reducing the Bcl-2 (p<0.01). Treatment with *L. rhamnosu*s tended to lower Bax and Caspase-3 expression and elevate Bcl-2 (Figure 3C). Western blot analyses of these oxidative and apoptotic proteins were consistent with the RT-PCR findings (Figure 3A–3B, 3D, 3F).

**Figure 2.**
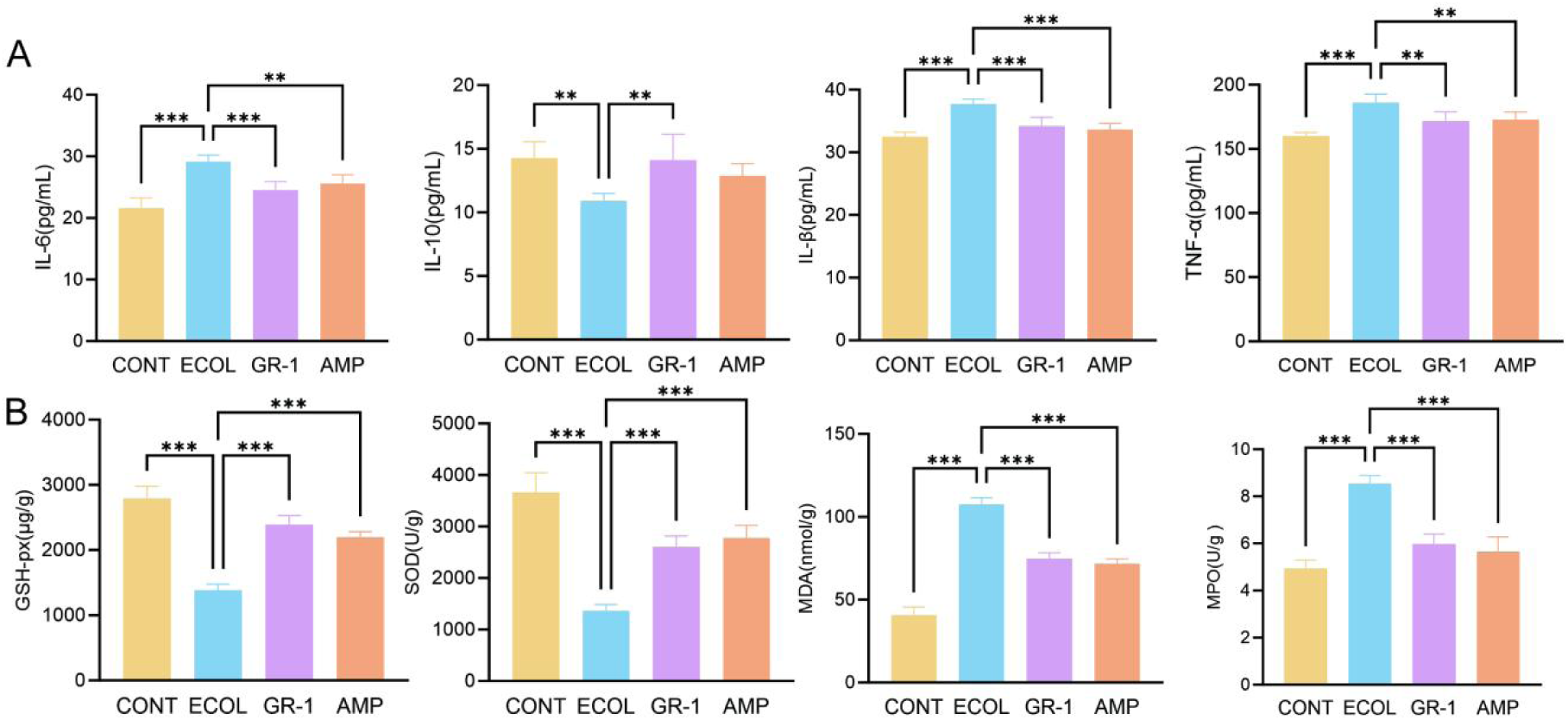
Effects of *L. rhamnosus* GR-1 on inflammation and oxidative stress levels in uterine tissue of mice induced by *E. coli.* (A) Relative expression levels of IL-6, IL-1β, IL-10, and TNF-α in mouse uterus detected by ELISA. (B) Contents of GSH, SOD, MDA, and MPO in mouse uterine tissue. Data represented are mean ± SD, *p<0.05; **p<0.01; ***p<0.001, n=6.

**Figure 3.**
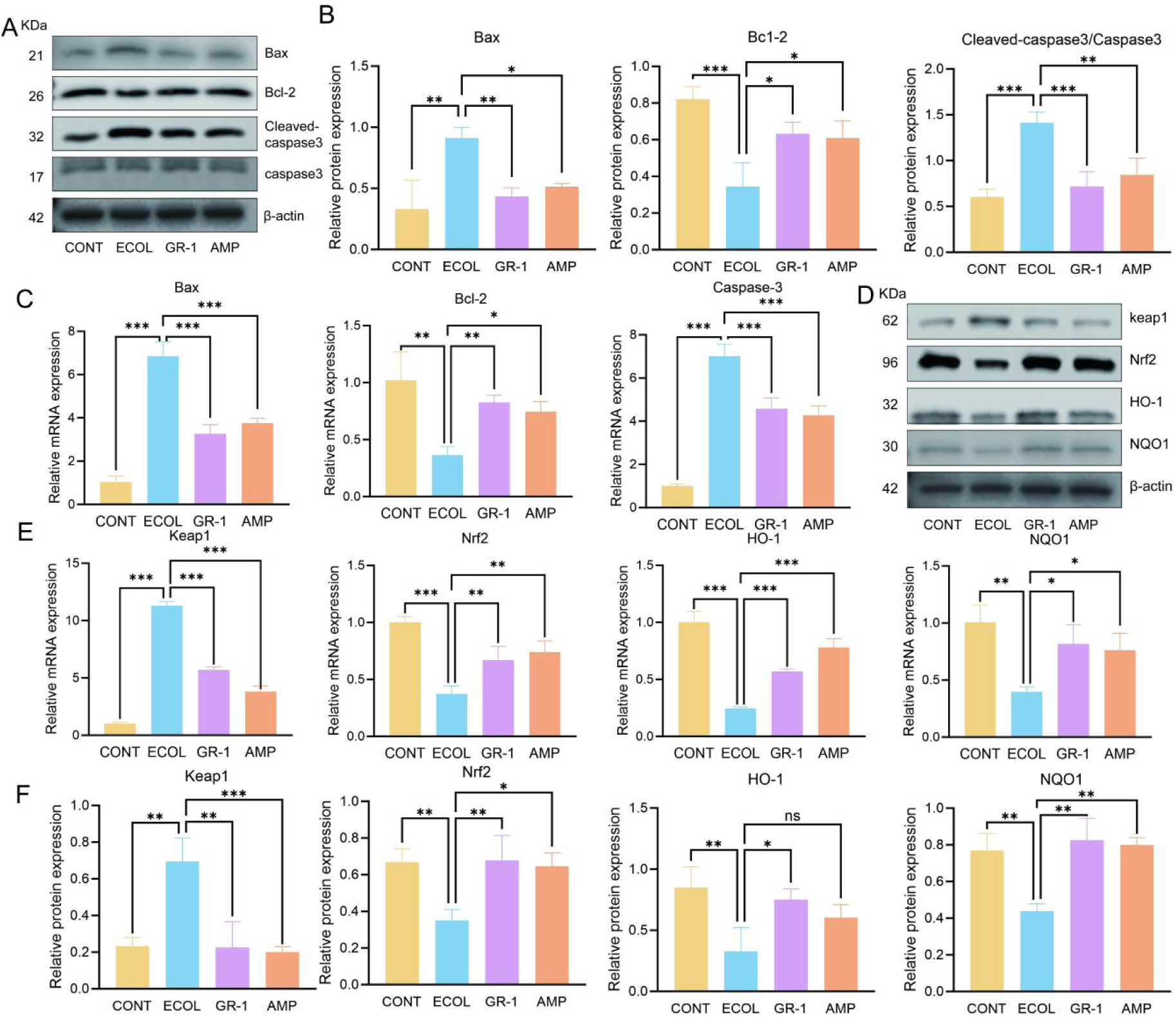
Effects of *L. rhamnosus* GR-1 on oxidation and apoptosis of mouse uterine tissue induced by *E. coli*. (A) Protein grayscale values of Bcl-2, Bax, Cleaved-caspase3, Caspase-3, and β-actin (internal standard). (B) Quantitative analysis of Bcl-2, Bax, and Cleaved-caspase3/Caspase-3 expression levels in three independent experiments, standardized using β-actin as the internal standard. (C) Relative mRNA expression levels of Caspase-3, Bcl-2, and Bax. (D) Protein grayscale values of HO-1, Nrf2, Keap1, NQO1, and β-actin (internal standard). (E) Relative mRNA expression levels of HO-1, Nrf2, Keap1, and NQO1. (F) Quantitative analysis of HO-1, Nrf2, Keap1, and NQO1 expression levels in three independent experiments, standardized using β-actin as the internal standard. Data represented are mean ± SD, *p<0.05; **p<0.01; ***p<0.001, n=3.

### Regulation of *L. rhamnosus* GR-1 on the uterine microflora of endometritis mice

Principal Coordinate Analysis (PCoA) and non-metric multidimensional scaling (NMDS) were performed using the Bray–Curtis dissimilarity matrix. As shown in Figure 4A–4B, PC1 and PC2 accounted for 26.76% and 18.39% of the overall variance, respectively, and the NMDS solution yielded a stress value of 0.08. The ECOL group contained 489 OTUs unique to that group, whereas the GR-1 group had 5308 unique OTUs, with 363 OTUs shared by ECOL and GR-1 (Figure 4C). Notable differences in microbial evenness and dominance were observed between the CONT and GR-1 groups (Figure 4D–4F). At the genus taxonomic level, *Rodentibacter* and *Escherichia–Shigella* predominated in the ECOL group (Figure 4G). At the phylum level, no single known taxon was exclusively dominant in either the GR-1 or AMP group; nonetheless, the GR-1 group showed higher representation of *Lactobacillales* and *Bacteroidales*, whereas *Enterobacteriales* remained relatively abundant in the AMP group (35%) (Figure 4H). Moreover, *Cutibacterium*, *Neisseria*, *Actinomyces*, *Rothia*, *Kocuria*, *Micrococcus*, and *Porphyromonas* were significantly enriched in the GR-1 group compared with the AMP group (Supplement 5). LEfSe analysis indicated that the LDA scores of discriminative taxa among groups were above 5 (Figure 4I–4J). *Lactobacillales*, *Propionibacteriales*, and *Bacteroidales* were characterized by higher LDA scores and richer abundances in the GR-1 group. The ECOL group exhibited significantly increased abundances of *Enterobacterales*, *Gammaproteobacteria*, and *Proteobacteria*.

**Figure 4.**
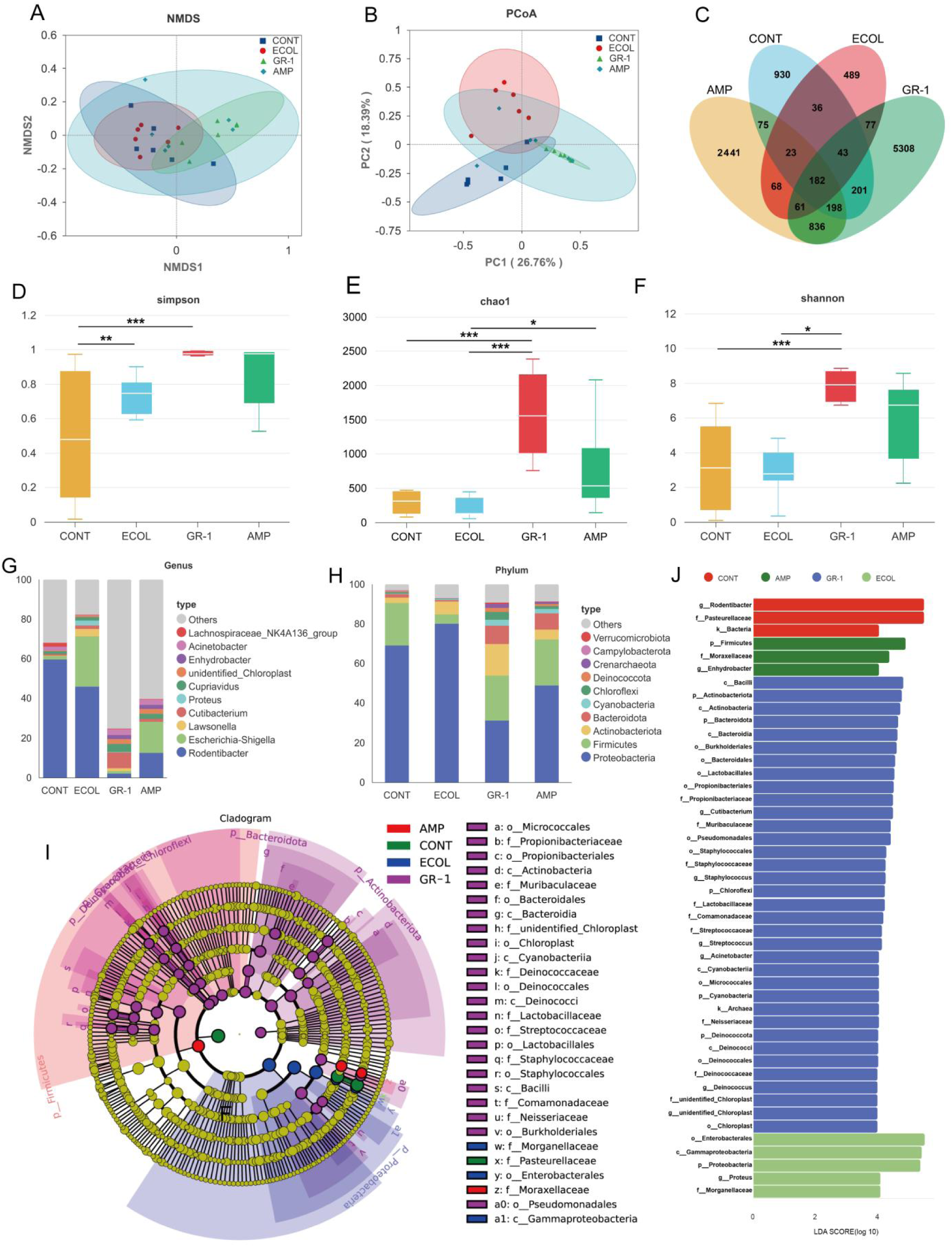
Analysis of Uterine Microbial Diversity and Community Structure Based on 16S rRNA Sequencing. (A) Non-metric Multidimensional Scaling (NMDS) analysis based on the Bray-Curtis metric. (B) Principal Coordinate Analysis (PCoA). (C) Analysis results of operational taxonomic units (OTU). (D-F) Analysis of differences in species diversity among the four groups of uterine microbiota in terms of chao1, Shannon, and Simpson indices. (G) Bacterial classification characteristics of the four groups of uterine microbiota at the genus level and (H) phylum level. (I) Evaluation of the degree of influence of changes in uterine microbiota among the four groups through the integration of Effect Size (LEfSe) and (J) Linear Discriminant Analysis (LDA), *p<0.05; **p<0.01; ***p<0.001, n=6.

### Regulatory effect of *L. rhamnosus* GR-1 on uterine metabolism in endometritis mice

A total of 148 differential metabolites were identified between the ECOL and GR-1 groups; 229 unique differential metabolites between the ECOL and AMP groups (Figure 5A). Among all annotated compounds, lipids and lipid-like molecules constituted the largest metabolite class, followed by organic acids and their derivatives, whereas categories such as organic heterocyclic compounds and organic oxygen-containing compounds accounted for smaller fractions (Figure 5B). Metabolomic profiling of uterine samples demonstrated clear metabolic separation between the ECOL and GR-1 groups, while the distinction between ECOL and AMP was comparatively less apparent (Figure 5C, 5E). During the screening of differential metabolites, a large number of differential metabolites exhibited enhanced expression after treatment with *L. rhamnosus* (Figure 5D, 5F). To interpret the biological relevance of these metabolic alterations, KEGG pathway enrichment analysis was conducted. Metabolites altered in the GR-1 group were significantly enriched in pathways including circadian rhythm, aldosterone-regulated sodium reabsorption, and neuroactive ligand–receptor interaction. In the AMP group, enriched pathways mainly involved bile secretion, glycine/serine/threonine metabolism, and lysine degradation (Figure 5G-5H).

**Figure 5.**
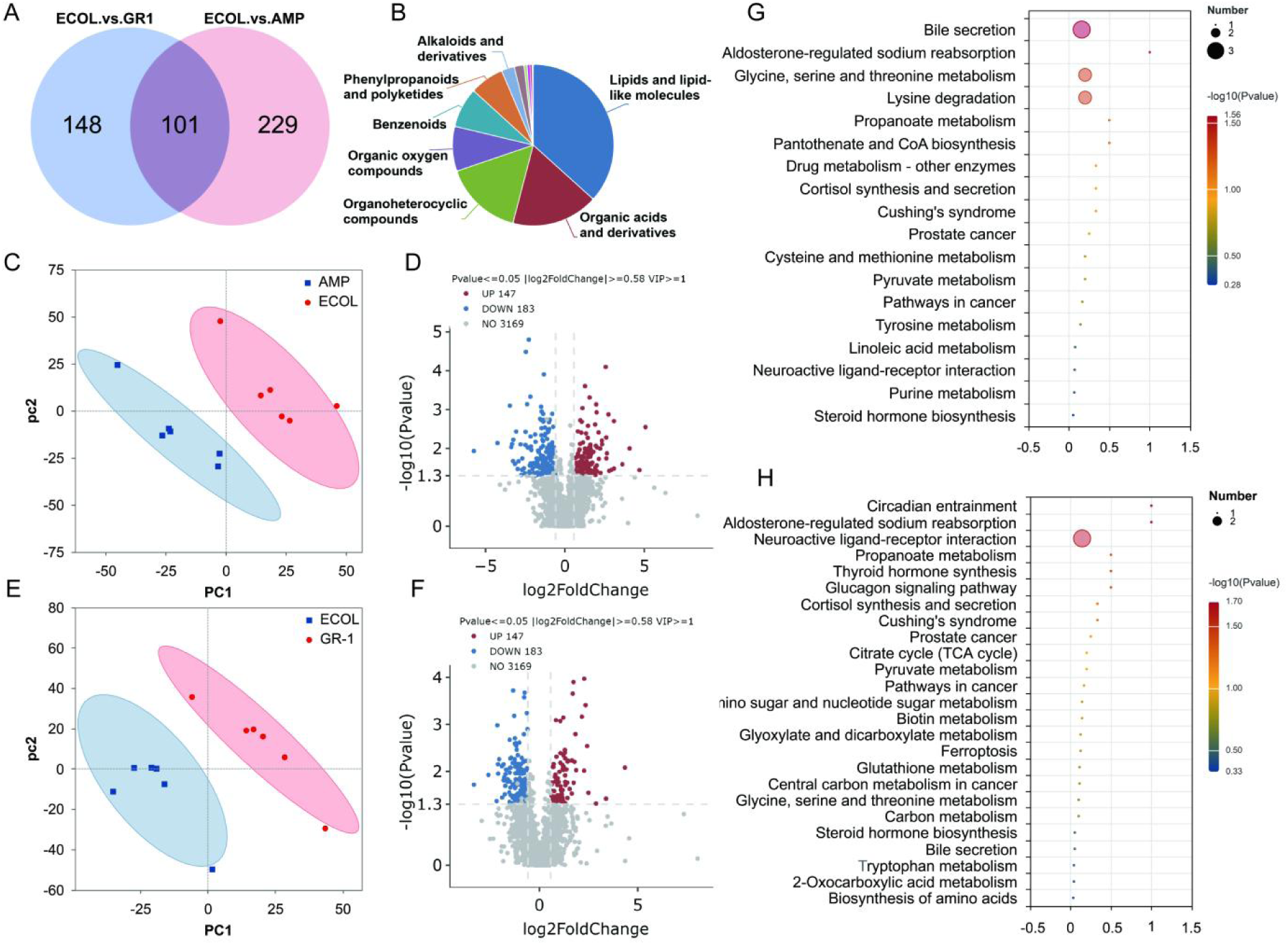
Multi-omics Difference. (A) Venn diagram of overlapping and group-specific differential metabolites. (B) Distribution of all metabolites. Distribution of differential metabolites: (C) ECOL group vs. AMP group and (E) ECOL group vs. GR-1 group. Expression levels of different differential metabolites: (D) ECOL group vs. AMP group and (F) ECOL group vs. GR-1 group. KEGG functional annotation and enrichment analysis of differential metabolites: (G) ECOL group vs. AMP group and (H) ECOL group vs. GR-1 group, n=6.

### Correlation analysis

To elucidate how probiotic treatment mitigates murine endometritis, we examined associations between uterine metabolites and bacterial genera (Figure 6A). All 30 metabolites included in the analysis showed significant correlations with at least one genus. Specifically, 7 metabolites displayed positive associations, whereas 23 metabolites were negatively associated. In addition, 23 metabolites were positively correlated with *Escherichia–Shigella* and *Rhodobacter*, while 7 metabolites showed significant negative correlations with these taxa. To further characterize microbiota–metabolite interactions in the ECOL versus GR-1 groups, we assessed correlations between differential metabolites and differential genera identified in these two groups. *Bacillus*, *Brevibacillus*, and *Deinococcus* were positively associated with L-isoleucyl-L-proline, N-acetylcysteine, and 10,16-dihydroxyhexadecanoic acid. In contrast, *Escherichia–Shigella* was negatively correlated with L-isoleucyl-L-proline and bebeerine. N-lactoyl-methionine showed positive correlations with the majority of microbial genera, whereas cytidine, oleoyltaurine, and oleoylethanolamide were negatively correlated with most microbial communities. We next compared uterine microbiota profiles with oxidative stress indices and inflammatory cytokines. *Escherichia–Shigella* and *Rodentibacter* were positively correlated with IL-6, IL-1β, TNF-α, MPO, and MDA. Conversely, *Bacillus* and *Brevibacillus* were negatively correlated with IL-6, IL-1β, and TNF-α, and positively correlated with IL-10, GSH, and SOD (Figure 6B).

**Figure 6.**
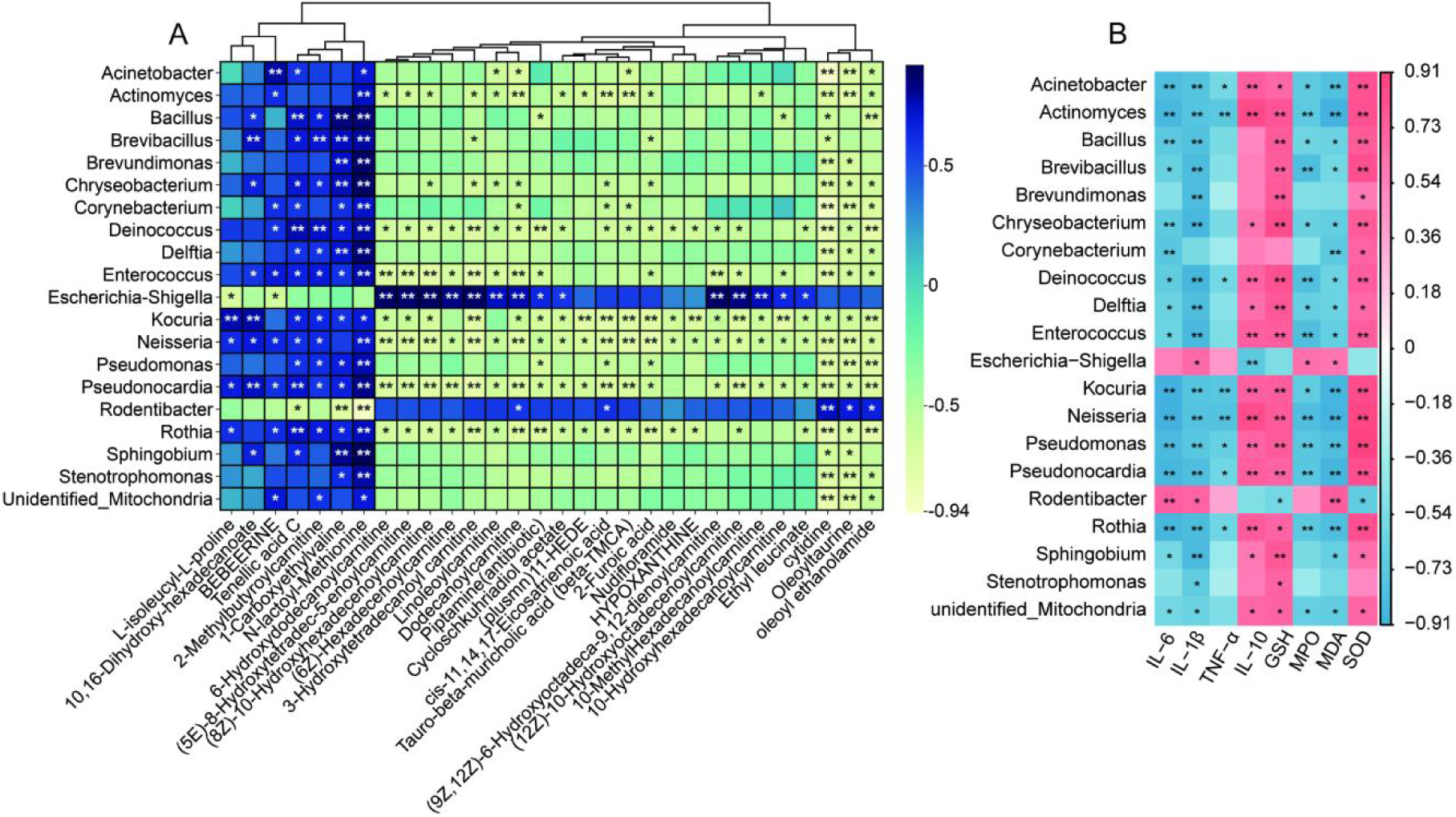
Correlation Analysis. (A) The Spearman analysis (heatmap) was used to analyze the correlation between differential metabolites and differential microorganisms between the ECOL group and the GR-1 group. (B) Spearman analysis (heat map) was performed to analyze the correlation between inflammatory markers, oxidative indicators, and changes in uterine microbiota. The color scale of the heat map indicates the Pearson correlation coefficients. In (A), darker blue and yellow represent higher absolute values, while in (B), darker red and blue indicate higher absolute values. *p<0.05; **p<0.01; ***p <0.001, n=6.

### Effects of *L. rhamnosus* GR-1 on reproductive index in endometritis mice

The primary harm caused by endometritis is the disruption of normal endometrial architecture and function, which in turn compromises embryo implantation and the maintenance of pregnancy (6). In the CONT group, the female pregnancy rate reached 100%, with an average litter size of 7.3 pups per dam and a mean total pup weight of 11.09 g. No mice became pregnant in the ECOL group. In the GR-1 group, the pregnancy rate was comparable to that of the CONT group; however, both the average number of pups per litter and the mean pup weight were lower than those observed in CONT. The AMP group showed reduced pregnancy rate, litter size, and pup weight compared with both the GR-1 and CONT groups. Notably, relative to the ECOL group, mice receiving GR-1 or AMP exhibited higher pregnancy rate, larger average litter size, and greater pup weight (Supplement 3).

## DISCUSSION

In dairy cows, bovine endometritis is characterized by inflammation of the uterine endometrium following childbirth [14]. *E. coli* induces persistent uterine infection and inflammation, particularly in postpartum or immunocompromised states [15]. Currently, the increasing prevalence of antimicrobial resistance underscore the urgent need for alternative therapeutic approaches [16]. Prior research has shown that *L. rhamnosus* GR-1 is capable of mitigating apoptosis in *E. coli*-infected bovine endometrial epithelial cells (BENDs) [17]. Our previous studies also confirmed that *L. rhamnosus* GR-1 exhibits inhibitory effects on *E. coli*-induced BENDs inflammatory damage [18]. However, its effect on endometritis has not been reported. Therefore, this study investigated the strategy of *L. rhamnosus* GR-1 in alleviating endometritis induced by *E. coli* infection in mice.

When dairy cows develop endometritis induced by *E. coli* infection, they typically present with elevated body temperature and some affected cows may exhibit signs of dehydration and emaciation [19]. In this study, following *E. coli* infection, mice displayed weight loss and elevated body temperature. This is the same as the clinical manifestations of endometritis caused by *E. coli* infection in cows. Furthermore, *L. rhamnosus* GR-1 alleviates inflammatory reactions by regulating the balance between proinflammatory and anti-inflammatory cytokines [18]. Our study revealed that *L. rhamnosus* GR-1 exerted a pronounced inhibitory effect on the expression of TNF-α, IL-1β, and IL-6, while enhancing the production of the IL-10. This is consistent with previous findings that the *L. rhamnosus* strain GR-1 is capable of mitigating the inflammation-mediated damage to BENDs challenged with *E. coli* [13]. Thus, *L. rhamnosus* GR-1 is capable of inhibiting the uterine inflammatory response induced by *E. coli* both. The surface proteins, extracellular polysaccharides, secreted short-chain fatty acids (SCFAs) and bacteriocins of *Lactobacillus* are the key factors to enhance the stability of the barrier structure and inhibit the proliferation of pathogens [20–22]. Our findings indicated that *L. rhamnosus* GR-1 can mitigate *E. coli* colonization, implying that the potential secretion of certain inhibitory compounds against *E. coli* by this bacterium.

Infection of dairy cows with *E. coli* triggers excessive production of ROS, leading to lipid peroxidation and cellular damage [23]. Nrf2 maintains its activity and stability by binding to Keap1 in the cytoplasm [24,25]. Our experimental results revealed that *L. rhamnosus* GR-1 intervention reversed *E. coli*-induced oxidative stress in the mouse uterus, including reduced Keap1 activity, increased SOD and GSH-Px activity, and enhanced Nrf2, HO-1, and NQO1 expression. This established an antioxidant network capable of scavenging excess ROS, thereby inhibiting *E. coli*-induced damage to uterine tissue. Following infection of dairy cow uterine epithelial cells with pathogenic *E. coli*, the bacteria directly induce nuclear damage phenotypes including chromosome condensation and abnormal nuclear morphology [15]. Meanwhile, the infection triggers excessive intracellular ROS production; excess ROS further exacerbates cellular damage, triggers the apoptosis-promoting protein Bax of the expression, activating caspase-3 sequentially, ultimately leading to cell apoptosis [18]. Studies have revealed that catalase modulates hepatic Bax expression, its cross-talk with Bcl-2, and diminishes caspase-3 and caspase-9 to alleviate oxidative damage in porcine liver tissue [26]. The present study confirmed that *L. rhamnosus* GR-1 intervention effectively reversed the abnormal expression of Bax, Cleaved-caspase3 and Bcl-2 in uterine tissues. *L. rhamnosus* GR-1 plays a key role in alleviating oxidative stress and apoptosis induced by *E. coli* in uterus.

In the treatment of endometritis, antibiotics remain the primary therapeutic approach. Ampicillin is one of the preferred options for managing endometritis in dairy cows [27]. However, ampicillin exert a certain disruptive effect on the balance of the uterine microecological environment [28]. In current study, the uterine microbiota of mice treated with ampicillin exhibited persistently high abundances of *Escherichia-Shigella*, such high levels of drug-resistant strains may account for the diminished therapeutic efficacy observed in ampicillin. Ampicillin markedly diminished the abundance of *Lactobacillaceae* and the *Lachnospiraceae* NK4A136, in the uterus while abnormally enriching conditional pathogens like *Comamonadaceae* and *Moraxellaceae*. Such dysbiosis compromises the uterus’s natural defense mechanisms and fosters an environment conducive to conditional pathogens, thereby increasing the risk of endometritis recurrence [29]. Prior research has revealed that *L. rhamnosus* GR-1 exerts modulatory effects on both vaginal and intestinal microbial communities [11], this finding that aligns with the results of our current investigation. The strain elevated the diversity of the uterine microbiota, reduced the abundance of pathogens such as *Enterobacteriaceae* and *Shigella*, and promoted the proliferation of *Lactobacillaceae* and *Bacteroidetes*, thereby facilitating the restoration of the microbial community’s balanced structure. This precisely compensates for the insufficient endometrial receptivity caused by persistent dysbiosis and metabolic disturbances that may arise from antibiotics, despite their effective control of inflammation [30], thereby creating a favorable uterine microbial environment for embryo implantation.

The dynamic equilibrium between microbiota and metabolism is crucial for maintaining homeostasis in the uterine microenvironment [31]. In the present investigation, the metabolite expression profiles of the GR-1 group differed significantly from those of the ECOL group. This difference may stem from the distinct mechanisms of action between probiotics and antibiotics. *L. rhamnosus* GR-1 regulated the uterine microecological balance by promoting the enrichment of bacterial populations such as *Bacillus* and *Bacillus brevispursus*. The “circadian entrainment” pathway is closely associated with inflammatory responses and tissue repair, enhancing cellular antioxidant capacity and damage repair efficiency [32]. Enrichment of this pathway implies that *L. rhamnosus* GR-1 might enhance the inflammation-suppressive and tissue-repairing capacity of the uterus via the modulation of circadian rhythms in uterine tissue cells. The aldosterone-regulated sodium reabsorption pathway plays a critical role in maintaining osmotic balance in uterine tissues. Activation of this pathway promotes sodium reabsorption, stabilizes the osmotic pressure of the uterine microenvironment, and alleviates tissue edema induced by infections [33]. This corresponds to the bacterium’s ability to reduce uterine tissue edema in mice caused by *E. coli* infection.

This study found that all 30 metabolites analyzed were significantly associated with at least one microbial genus, *Escherichia-Shigella* showed positive correlation with 23 metabolites, and *Escherichia* is a pathogenic genus that causes endometritis, indicating that these metabolites are closely related to inflammatory diseases. The negative correlation between some metabolites and *Escherichia spp.* May represent the host’s compensatory metabolic response to infection or the inhibitory effect of specific metabolites on pathogenic bacteria. N-acetylcysteine (NAC) can eliminate excess ROS generated by *E. coli* infection by regulating GSH-Px, thereby reducing oxidative stress damage [34]. As a hydroxy fatty acid derivative, 10,16-dihydroxyhexadecanoic acid exerts inflammation-alleviating effects via the inhibition of the NF-κB pathway [35]. *Bacillus, Brevibacillus and Deinococcus* exhibited positive correlations with L-isoleucyl-L-proline, NAC and 10,16-dihydroxyhexadecanoic acid, implying that *L. rhamnosus* GR-1 may exert inflammation-suppressive and antioxidant effects by enriching these bacterial taxa. As an alkaloid, the research has confirmed that bebeerine can inhibit the growth of Gram-negative bacteria [36]. In this study, *Escherichia* species showed negative correlations with L-isobutyryl-L-proline and bebeerine, suggesting that the reduction of *Escherichia* species in the GR-1 group may be partly attributed to the accumulation of these antimicrobial metabolites. Metabolic stress triggers inflammatory responses, which subsequently disrupt metabolic homeostasis [37]. *E. coli* and *Rodentibacter* induce excessive inflammatory responses and oxidative stress by secreting virulence factors such as lipopolysaccharide, leading to uterine tissue damage [38]. *E. coli* and *Rodentibacter* positively correlate with pro-inflammatory cytokines IL-6, IL-1β, TNF-4, MPO, MDA and negatively with IL-10, GSH, SOD, this aligns with our results and reflects their adverse effects on uterine tissue. In stark contrast, the genera *Bacillus* and *Brevibacillus* exhibited negative correlations with pro-inflammatory cytokines and positive correlations with IL-10 and antioxidant enzymes. This verified that *L. rhamnosus* GR-1 maight suppress host uterine inflammation by enriching this microbial community.

In this study, typical pathological changes of endometritis, including edema, inflammatory cell infiltration, and structural disorder, were observed in the uteri of mice after *E. coli* infection, and no pregnancy occurred in these mice. This indicates that endometritis caused by *E. coli* infection exerts an adverse effect on embryo implantation. After ampicillin treatment, although uterine inflammation was alleviated to a certain extent, the pregnancy rate and litter size of mice could not be restored. Consistent with previous research results, antibiotics can alleviate *E. coli*-induced endometritis but have poor effects on restoring reproductive performance [39], indicating the limitations of antibiotic therapy. Studies have shown that *Clostridium butyricum* can improve the reproductive performance of mice with *E. coli*-induced endometritis [6]. In line with our findings, this indicated that the strain could improve the reproductive capacity of mice suffering from endometritis. This highlights the potential of *L. rhamnosus* GR-1 in achieving inflammation resolution and improving the litter-bearing capability of mice. This study evaluated the therapeutic efficacy of *L. rhamnosus* GR-1 in an *E. coli*–induced mouse endometritis model and described its associations with uterine microbiota and metabolite changes; however, the causal mechanisms of the uterine microbiota–metabolite axis were not determined. Validation in a dairy cow endometritis model is warranted to assess translational efficacy.

## CONCLUSION

In summary, *L. rhamnosus* GR-1 alleviated *E. coli* – induced endometritis in mice by enhancing anti-inflammatory and antioxidant defenses and suppressing oxidative apoptosis. These effects were accompanied by favorable shifts in the uterine microbiota and metabolite profiles and improved pregnancy outcomes, suggesting GR-1 as a promising antibiotic alternative for endometritis. Further validation in dairy cows is warranted.

## AUTHORS’ CONTRIBUTION

Conceptualization: Li X, Jia L.

Data curation: Li X, Wang M, Jia L.

Formal analysis: Wen X, Wang M.

Methodology: Wen X, Song D.

Software: Jia L, Li X, Song D.

Validation: Liang G, Liu D.

Investigation: Li X, Liu M.

Writing – original draft: Li X, Jia L.

Writing – review & editing: Li X, Jia L, Wang M, Wen X, Song D, Liang G, Liu D, Liu M.

## ACKNOWLEDGMENTS

The authors want to express their sincere gratitude to the members of the laboratory for their professional technical support.

## SUPPLEMENTARY MATERIAL

Supplement 1. Uterine injury assessment criteria.

Supplement 2. Primers used for the qRT-PCR study.

Supplement 3. Statistics of mouse reproduction.

Supplement 4. Effects of *L. rhamnosus* GR-1 on the uterus of mice.

Supplement 5. The microbiota correlation network of the GR-1 group and the AMP group.

## DATA AVAILABILITY

The dataset used in this study is available from the corresponding author upon reasonable request and has also been submitted to the National Center for Biotechnology Information with the accession number PRJNA1344922.

## ETHICS APPROVAL

All procedures involving mice followed the guidelines approved by the Animal Care and Use Committee of Hebei Agricultural University and were approved by the university’s Animal Ethics Committee(Approval Number: 2021041).

## DECLARATION OF GENERATIVE AI

No AI tools were used in this article.

## REFERENCES

1. Yang X, Zhang S, Liu B, et al. Exogenous Prostaglandin D(2) as a Modulator in Bovine Endometritis: Implications for Reducing Antibiotic Use in Dairy Cattle. Front Vet Sci 2025;12:1618203. 10.3389/fvets.2025.1618203

2. Gonzalez Moreno C, Torres Luque A, Oliszewski R, Rosa RJ, Otero MC. Characterization of Native Escherichia Coli Populations from Bovine Vagina of Healthy Heifers and Cows with Postpartum Uterine Disease. PloS one 2020;15:e0228294. 10.1371/journal.pone.0228294

3. He Y, Cai J, Xie X, et al. Dimethyl Itaconate Alleviates *Escherichia Coli*-Induced Endometritis through the Guanosine-Cxcl14 Axis Via Increasing the Abundance of Norank_F_Muribaculaceae. Adv Sci (Weinh) 2025;12:e2414792. 10.1002/advs.202414792

4. Xiao J, Li S, Zhang R, et al. Isgylation Inhibits an Lps-Induced Inflammatory Response Via the Tlr4/Nf-Κb Signaling Pathway in Goat Endometrial Epithelial Cells. Animals (Basel) 2021;11:2593. 10.3390/ani11092593

5. Ma X, Yin B, Guo S, et al. Enhanced Expression of Mir-34a Enhances *Escherichia Coli* Lipopolysaccharide-Mediated Endometritis by Targeting Lgr4 to Activate the Nf-Κb Pathway. Oxid Med Cell Longev 2021;2021:1744754. 10.1155/2021/1744754

6. Mun C, Cai J, Hu X, Zhang W, Zhang N, Cao Y. Clostridium Butyricum and Its Culture Supernatant Alleviate the *Escherichia Coli*-Induced Endometritis in Mice. Animals (Basel) 2022;12:2719. 10.3390/ani12192719

7. Jiang K, Yang J, Song C, He F, Yang L, Li X. Enforced Expression of Mir-92b Blunts E. Coli Lipopolysaccharide-Mediated Inflammatory Injury by Activating the Pi3k/Akt/Β-Catenin Pathway Via Targeting Pten. Int J Biol Sci 2021;17:1289–301. 10.7150/ijbs.56933

8. Zhang K, Feng, H, Zhang J, et al. Prevalence and Molecular Characterization of Extended-Spectrum Β-Lactamase-Producing *Escherichia Coli* Isolates from Dairy Cattle with Endometritis in Gansu Province, China. BMC Vet Res 2024;20:19. 10.1186/s12917-023-03868-x

9. Dias NW. Editorial: Understanding the Female Reproductive Microbiome in Livestock. Front Microbiol 2025;16:1570990. 10.3389/fmicb.2025.1570990

10. Su Z, Tian C, Wang G, Guo J, Yang X. Study of the Effect of Intestinal Microbes on Obesity: A Bibliometric Analysis. Nutrients 2023;15:3255. 10.3390/nu15143255

11. Borum LS, Bartolomaeus TUP, Lamont RF, et al. Probiotic Ice Cream Influences Gut and Vaginal Microbiota in Women at High Risk of Preterm Birth: A Randomized Controlled Study. Matern Health Neonatol Perinatol 2025;11:43. 10.1186/s40748-025-00238-3

12. Martínez JE, Vargas A, Pérez-Sánchez T, Encío IJ, Cabello-Olmo M, Barajas M. Human Microbiota Network: Unveiling Potential Crosstalk between the Different Microbiota Ecosystems and Their Role in Health and Disease. Nutrients 2021;13:2905. 10.3390/nu13092905

13. Feng X, Li Y, Wen X, et al. *Lactobacillus Rhamnosus* Gr-1 Alleviates *Escherichia Coli*-Triggered Bovine Endometrial Epithelial Cells Damage Via the Reactive Oxygen Species-Mitochondrial Pathway. Anim Biosci 2025;38:1996–2007. 10.5713/ab.25.0031

14. Dong J, Ji B, Jiang Y, et al. A20 Alleviates the Inflammatory Response in Bovine Endometrial Epithelial Cells by Promoting Autophagy. Animals (Basel) 2024;14:2876. 10.3390/ani14192876

15. Wei S, Ding B, Wang G, Luo S, Zhao H, Dan X. Population Characteristics of Pathogenic *Escherichia Coli* in Puerperal Metritis of Dairy Cows in Ningxia Region of China: A Systemic Taxa Distribution of Virulence Factors and Drug Resistance Genes. Front Microbiol 2024;15:1364373. 10.3389/fmicb.2024.1364373

16. Lu Y, Geng W, Li L, et al. Enhanced Antibacterial and Antibiofilm Activities of Quaternized Ultra-Highly Deacetylated Chitosan against Multidrug-Resistant Bacteria. Int J Biol Macromol 2025;298:140052. 10.1016/j.ijbiomac.2025.140052

17. Lai JL, Liu YH, Liu C, et al. Indirubin Inhibits Lps-Induced Inflammation Via Tlr4 Abrogation Mediated by the Nf-Kb and Mapk Signaling Pathways. Inflammation 2017;40:1–12. 10.1007/s10753-016-0447-7

18. Liu J, Feng X, Li B, et al. *Lactobacillus Rhamnosus* Gr-1 Alleviates *Escherichia Coli*-Induced Inflammation Via Nf-Κb and Mapks Signaling in Bovine Endometrial Epithelial Cells. Front Cell Infect Microbiol 2022;12:809674. 10.3389/fcimb.2022.809674

19. Chirivi M, Contreras GA. Endotoxin-Induced Alterations of Adipose Tissue Function: A Pathway to Bovine Metabolic Stress. J Anim Sci Biotechnol 2024;15:53. 10.1186/s40104-024-01013-8

20. Patki A, Kar S, Patel N, Ingale K, Bansal K, Durga P. Expert Opinion: Place in Therapy of Probiotics in Infertility and Recurrent Implantation Failure. Cureus 2025;17:e81067. 10.7759/cureus.81067

21. Matsuzaki C, Takagi H, Saiga S, et al. Prebiotic Effect of Galacto-N-Biose on the Intestinal Lactic Acid Bacteria as Enhancer of Acetate Production and Hypothetical Colonization. Appl Environ Microbiol 2024;90:e0144523. 10.1128/aem.01445-23

22. Skoufou M, Tsigalou C, Vradelis S, Bezirtzoglou E. The Networked Interaction between Probiotics and Intestine in Health and Disease: A Promising Success Story. Microorganisms 2024;12:194. 10.3390/microorganisms12010194

23. Darfarin G, Pluth J. Mitochondria-Nuclear Crosstalk: Orchestrating Mtdna Maintenance. Environ Mol Mutagen 2025;66:222–42. 10.1002/em.70013

24. Zhang N, Wang W, Zhang R, et al. Melatonin Alleviates Oral Epithelial Cell Inflammation Via Keap1/Nrf2 Signaling. Int J Immunopathol Pharmacol 2025;39:3946320251318147. 10.1177/03946320251318147.

25. Jiang Y, He P, Sheng K, et al. The Protective Roles of Eugenol on Type 1 Diabetes Mellitus through Nrf2-Mediated Oxidative Stress Pathway. ELife 2025;13:RP96600. 10.7554/eLife.96600

26. Wang W, Zhu J, Cao Q, et al. Dietary Catalase Supplementation Alleviates Deoxynivalenol-Induced Oxidative Stress and Gut Microbiota Dysbiosis in Broiler Chickens. Toxins 2022;14:830. 10.3390/toxins14120830

27. Su J, Yang H, Wang S, et al. An Engineered Antimicrobial Peptide Z-Fv7 Demonstrates Bactericidal Efficacy against Multidrug-Resistant *Escherichia Coli* in a Murine Model of Endometritis. Vet Res 2025;56:159. 10.1186/s13567-025-01593-x

28. Nisansala T, Gunasekara YD, Piyarathne NS. Phenotypic and Genotypic Landscape of Antibiotic Resistance through One Health Approach in Sri Lanka: A Systematic Review. Trop Med Int Health 2025;30:143–58. 10.1111/tmi.14084

29. Kim MJ, Yoo J, Yoo S, Kwon MY, Lee S, Kim M. The Clinically Significant Changes in the Composition and Functional Diversity of the Vaginal Microbiome in Women with Type 2 Diabetes Mellitus. Microorganisms 2025;13:1426. 10.3390/microorganisms13061426

30. Guo H, Chen L, Liu L. Study on the Effects of Doxycycline Treatment on Endometrial Microbiota and Pregnancy Outcomes in Chronic Endometritis. Am J Reprod Immunol 2025;94:e70184. 10.1111/aji.70184

31. Larnder AH, Manges AR, Murphy RA. The Estrobolome: Estrogen-Metabolizing Pathways of the Gut Microbiome and Their Relation to Breast Cancer. Int J Cancer 2025;157:599–613. 10.1002/ijc.35427

32. Zhang H, Li K, Wang Y, et al. Ros Regulates Circadian Rhythms by Modulating Ezh2 Interactions with Clock Proteins. Redox Biol 2025;81:103526. 10.1016/j.redox.2025.103526

33. Lv L, Mu D, Du Y, Yan R, Jiang H. Mechanism of the Immunomodulatory Effect of the Combination of Live Bifidobacterium, Lactobacillus, Enterococcus, and Bacillus on Immunocompromised Rats. Front Immunol 2021;12:694344. 10.3389/fimmu.2021.694344

34. Ni D, Tang T, Lu Y, et al. Canonical Secretomes, Innate Immune Caspase-1-, 4/11-Gasdermin D Non-Canonical Secretomes and Exosomes May Contribute to Maintain Treg-Ness for Treg Immunosuppression, Tissue Repair and Modulate Anti-Tumor Immunity Via Ros Pathways. Front Immunol 2021;12:678201. 10.3389/fimmu.2021.678201

35. Wang Z, Yin Y, Mu Y, et al. Exploring the Occurrence Mechanism and Early-Warning Model of Phlebitis Induced by Aescinate Based on Metabolomics in Cerebral Infarction Patients. J Inflamm Res 2024;17:343–55. 10.2147/jir.S436846

36. Kurnia D, Lestari S, Mayanti T, Gartika M, Nurdin D. Anti-Infection of Oral Microorganisms from Herbal Medicine of Piper Crocatum Ruiz & Pav. Drug Des Devel Ther 2024;18:2531–53. 10.2147/dddt.S453375

37. Lu M, Li W, Zhou J, et al. Integrative Bioinformatics Analysis for Identifying the Mitochondrial-Related Gene Signature Associated with Immune Infiltration in Premature Ovarian Insufficiency. BMC Med 2024;22:444. 10.1186/s12916-024-03675-7

38. Yang X, Li X, Guo L, et al. Investigating the Role of the Mpges-Pge₂-Ep4 Pathway in *Escherichia Coli*-Induced Mastitis in Dairy Cows: Insights for Non-Antibiotic Therapeutic Strategies. Front Vet Sci 2025;12:1628028. 10.3389/fvets.2025.1628028

39. Bruinjé TC, Morrison EI, Ribeiro ES, Renaud DL, Couto Serrenho R, LeBlanc SJ. Postpartum, Health. Is Associated with Detection of Estrus by Activity Monitors and Reproductive Performance in Dairy Cows. J Dairy Sci 2023;106:9451–73. 10.3168/jds.2023-23268.

