## Supplementary Material for "*Lacticaseibacillus rhamnosus* GR-1 attenuates *Escherichia coli*-induced endometritis in mice, accompanied by modulation of uterine microbiota and metabolite profiles"

**Supplement 1. Uterine injury assessment criteria**

| Uterine endometrial injury | | Inflammatory infiltration | | Uterine edema | | Endometrial thickness | |
| --- | --- | --- | --- | --- | --- | --- | --- |
| Grade | Score | Grade | Score | Grade | Score | Grade | Score |
| Normal | **0** | Normal | **0** | Normal | **0** | Normal | **0** |
| + | **1** | **+** | **1** | **+** | **1** | **+** | **1** |
| **++** | **2** | **++** | **2** | **++** | **2** | **++** | **2** |
| **+++** | **3** | **+++** | **3** | **+++** | **3** | **+++** | **3** |

The "+" symbol indicates the degree of uterine injury.

**Supplement 2.** Primers used for the qRT-PCR study

| **Gene** |  | **Primer (5'to 3')** |
| --- | --- | --- |
| ***GAPDH*** | F: | AGCTTGTCATCAACGGGAAG |
|  | R: | TTTGATGTTAGTGGGGTCTCG |
| ***IL-1β*** | F: | GGGCCTCAAAGGAAAGAATC |
|  | R: | TACCAGTTGGGGAACTCTGC |
| ***IL-6*** | F: | CAAAGCCAGAGTCCTTCAGAG |
|  | R: | GCCACTCCTTCTGTGACTCC |
| ***TNF-α*** | F: | CAGGCGGTGCCTATGTCTC |
|  | R: | CGATCACCCCGAAGTTCAGTAG |
| ***Caspase-3*** | F: | TCTGACTGGAAAGCCGAAACTCTTC |
|  | R: | GTCCCACTGTCTGTCTCAATGCC |
| ***Bax*** | F: | TGCAGAGGATGATTGCTGAC |
|  | R: | GATCAGCTCGGGCACTTTAG |
| ***Bcl-2*** | F: | CCAGCCTGAGAGCAACCCAATG |
|  | R: | ACGACGGTAGCGACGAGAGAAG |
| ***NQO-1*** | F: | AGCCAATCAGCGTTCGGTAT |
|  | R: | GCCTCCTTCATGGCGTAGTT |
| ***Nrf2*** | F: | AAAATCATTAACCTCCCTGTTGAT |
|  | R: | CGGCGACTTTATTCTTACCTCTC |
| ***Keap1*** | F: | GATATGAGCCAGAGCGGGAC |
|  | R: | CATACAGCAAGCGGTTGAGC |
| ***HO-1*** | F: | CAAGCCGAGAATGCTGAGTTCATG |
|  | R: | GCAAGGGATGATTTCCTGCCAG |

**Supplement** **3. Statistics of mouse reproduction**

| Item | CONT | ECOL | GR-1 | AMP |
| --- | --- | --- | --- | --- |
| Number of mice in the experiment | 3 | 3 | 3 | 3 |
| Number of pregnant mice | 3 | 0 | 3 | 1 |
| conception rate(%) | 100 | 0 | 100 | 33.3 |
| Number of young per nest | 7 | 0 | 6 | 6 |
|  | 8 | 0 | 6 | 0 |
|  | 7 | 0 | 5 | 0 |
| Overall weight of pups (g) | 10.91 | 0 | 10.24 | 8.57 |
|  | 11.62 | 0 | 9.86 | 0 |
|  | 10.73 | 0 | 9.65 | 0 |

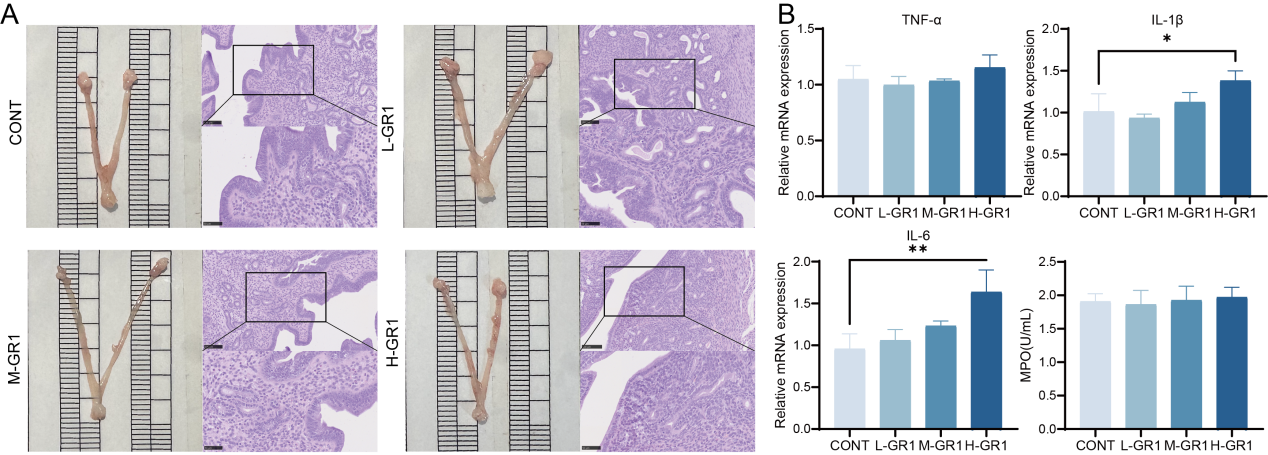

**Supplement** **4.** Effects of *L. rhamnosus* GR-1 on the uterus of mice. (A) Changes in uterine morphology and histopathology.(B) Changes in inflammation-related indicators (IL-1β, IL-6, TNF-α, and MPO) in uterine tissue. Groups: CONT (PBS), L-GR1 (*L. rhamnosus* GR-1, 1.0×10⁸ CFU), M-GR1 (*L. rhamnosus* GR-1, 1.0×10⁹ CFU), H-GR1 (*L. rhamnosus* GR-1, 1.0×10¹⁰ CFU). Data are expressed as mean ± SE.**p*<0.05 and ***p*<0.01 compared with the CONT group (n=6). Mice in each group were given uterine perfusion with the corresponding treatment at a volume of 100 μL per day for 6 consecutive days. On the 7th day, the mice were euthanized. Afterwards, the morphological changes of the uterine tissue were observed, and the indicators in the mouse uterine tissue were detected.

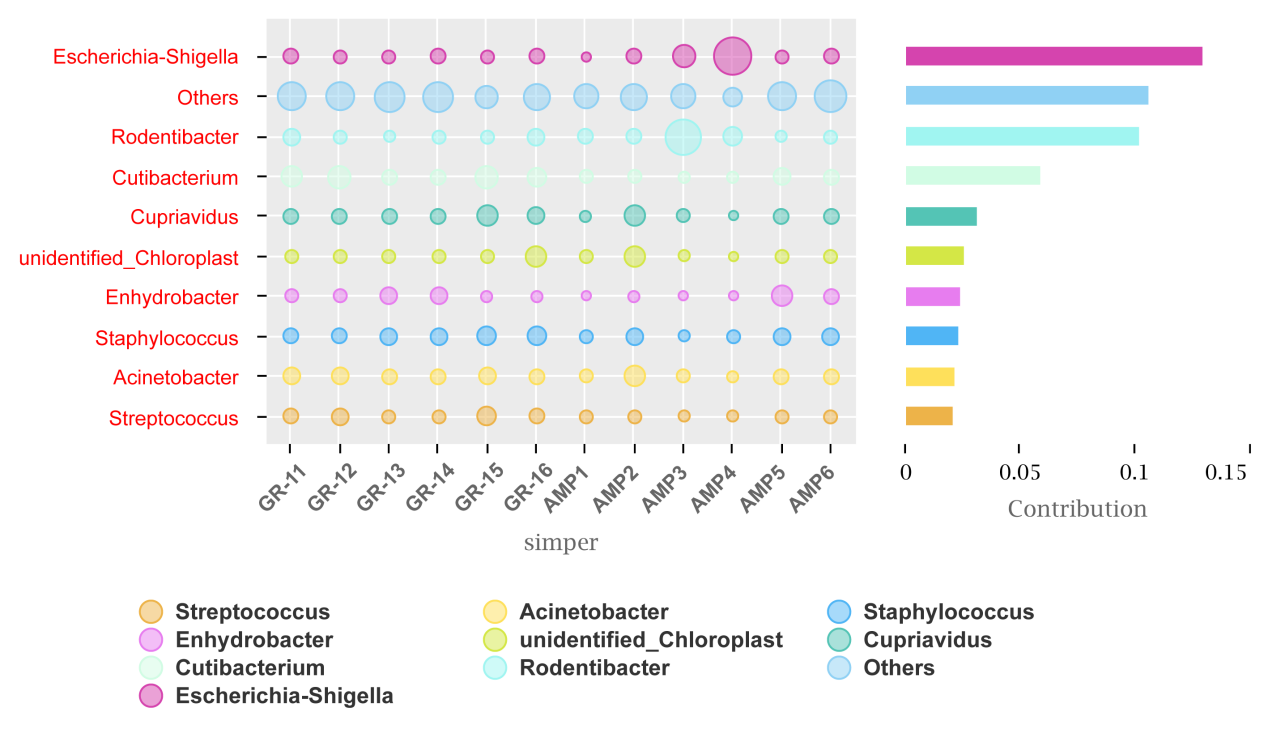

**Supplement** **5.** The microbiota correlation network of the GR-1 group and the AMP group. The size of the bubbles represents the proportion of the contribution of the corresponding microorganism in the respective samples.
